# An artefact-resilient wide bandwidth bidirectional graphene neural interface

**DOI:** 10.1101/2025.05.15.653988

**Authors:** Michał Prokop, Martín Esparza-Iaizzo, Eduard Masvidal-Codina, Xavi Illa, Neela K. Codadu, Daman Rathore, Nicola Ria, Kostas Kostarelos, Elena del Corro, Ramon Garcia-Cortadella, Rob C. Wykes, Anton Guimerà-Brunet, Jose A. Garrido

**Affiliations:** Catalan Institute of Nanoscience and Nanotechnology (ICN2), CSIC and BIST, Campus UAB, Barcelona, Spain; University College London, Queen Square Institute of Neurology, London, UK; INBRAIN Neuroelectronics SL, Barcelona, Spain; Institut de Microelectrònica de Barcelona, IMB-CNM (CSIC), Campus UAB, Bellaterra, Spain; Centro de Investigación Biomédica en Red de Bioingeniería, Biomateriales y Nanomedicina, Instituto de Salud Carlos III, Madrid, Spain; Institució Catalana de Recerca i Estudis Avançats (ICREA), Barcelona, Spain; Institute of Neurosciences, UAB Medical School, Bellaterra, Spain; Bernstein Center for Computational Neuroscience Munich, Faculty of Medicine, Ludwig-Maximilians Universität München, Planegg-Martinsried, Munich, Germany; Centre for Nanotechnology in Medicine & Division of Neuroscience, University of Manchester, Manchester, UK

## Abstract

The ability to simultaneously record and modulate neural activity is critical for next-generation bidirectional neural interfaces aiming to enable adaptive neuromodulation therapies for neurological disorders. Active graphene transistor technologies are particularly promising for neural recordings, as they extend the ability to monitor brain activity at very low frequencies and support multiplexed operation for high-density neural interfaces. However, their limited charge injection capacity makes them unsuitable for stimulation. In this work, we present a bidirectional neural interface that combines nanoporous reduced graphene oxide (rGO) microelectrodes for high charge injection focal stimulation and graphene solution-gated field-effect transistors (gSGFETs) for brain activity monitoring, exploiting the advantages of both technologies in one single device. Using scalable cleanroom microfabrication techniques, we monolithically integrate these two graphene-based technologies into fully flexible probes. We evaluate the performance of the hybrid devices both in saline and *in vivo*, with a particular focus on transistor performance during stimulation. Our results demonstrate that the recording capability of this bidirectional neural interface, including the monitoring of infraslow and local field potential activity, is not compromised during stimulation. This work highlights the potential of this hybrid neural interface for both basic neurophysiological and clinical translation use.

## Main

Brain implants have proven useful in treating common neurological disorders such as Parkinson’s disease (PD) [1,2,3], epilepsy [4,5], Tourette syndrome [6,7], essential tremor [8,9] and dystonia [10], among others. Most clinically available implants are unidirectional, meaning they can only electrically stimulate neural activity and lack the ability to adjust it based on dynamic changes in neural signals. Implantable neural interfaces that enable bidirectional communication, can allow recording and modulation of neural activity, and offer the potential for more effective treatments. These systems can provide on-demand, patient-specific, closed-loop therapy based on detected biomarkers, which is expected to minimize side-effects and extend device battery life [11,12,13,14]. A recent feasibility clinical study on PD treatment demonstrated that adaptive deep brain stimulation significantly improved motor symptoms and quality of life compared to the standard unidirectional stimulation [15].

At present, clinically available neural implants primarily use passive metallic electrodes for both recording and stimulation. However, these passive electrodes have inherent limitations, particularly in their ability to capture the full spectrum of brain activity with high spatial resolution. While passive electrodes can effectively record local field potentials (LFPs) (0.5– 200 Hz), high-frequency oscillations (HFOs) (200–600 Hz) and single spikes, they face challenges in detecting low-frequency signals such as DC-shifts and infraslow activity (ISA) (<0.1 Hz). This has been attributed to the instability of the metal-electrolyte interface, electrode potential drifts, and signal attenuation caused by high electrode impedance at low frequencies [16]. Despite these limitations, there is increasing interest in DC-coupled recordings due to mounting evidence linking ISA and DC-shifts with a range of neurological disorders, including epilepsy, stroke, migraine, and traumatic brain injury [17,18]. Moreover, recent evidence also links ISA and DC-shifts to physiological states that reflect varying levels of neural synchrony [19].

Graphene-based solution-gated field-effect transistors (gSGFETs) have emerged as a promising alternative to traditional passive electrodes, since they can offer high sensitivity for detecting ISA while also capturing high-frequency signals [20,21,22,23]. Among the various transistor technologies that have been validated for chronic in vivo electrophysiology [24,25], gSGFETs stand out due to the combination of graphene’s properties including electrochemical inertness, high carrier mobility, biocompatibility, and flexibility. Previous work has demonstrated that gSGFETs enable DC-coupled recordings and support multiplexed operation [26,27,28]. However, while gSGFETs are particularly suited for recording, they face limitations in stimulation because the low interfacial capacitance of single-layer graphene (typically 2 µF cm^-2^) restricts its charge injection capacity [29,30,31]. As a result, gSGFETs require integration with stimulating electrodes to enable wideband bidirectional communication with the nervous system.

To address the challenge of creating a bidirectional neural interface enabling closed-loop neuromodulation based on wideband recordings, we propose a novel device architecture that integrates gSGFETs for brain monitoring with passive reduced graphene oxide (rGO) microelectrodes for stimulation. Previous studies have highlighted the potential of rGO as a microelectrode material for high precision neural stimulation, owing to its low impedance and high charge injection [31,32,33]. The highly porous structure of rGO, which results in a high surface-to-volume ratio, enables it to deliver stimulation at high charge density even when miniaturized to the micrometer scale and integrated into flexible microfabricated arrays [31,32,33].

The integration of both recording and stimulation modalities within the same device introduces several technical challenges related to the bidirectional interface’s performance. Stimulation pulses can interfere with sensing instrumentation by generating artefacts that can obscure the underlying neural activity [34]. These artefacts, typically being several orders of magnitude larger than the biological signals of interest, can saturate the recording electronics. Understanding and mitigating the effect of stimulation artefacts on the recorded signals is critical for developing a new generation of closed-loop systems capable of providing on-demand stimulation [34].

Here, we present a wafer-scale microfabrication process that monolithically integrates gSGFETs with rGO microelectrodes, resulting in a fully flexible, ultra-thin neural probe. We have evaluated the bidirectional capability of this technology demonstrating that it can capture with high fidelity the high-amplitude stimulation-induced pulses without distortion. This ability facilitates the use of simple signal filtering techniques to recover brain signals below the stimulation frequency, enabling the simultaneous monitoring of low-amplitude local field potentials (LFPs) during electrical stimulation. Furthermore, we report a series of proof-of-concept *in vivo* studies that showcase the wideband recording capabilities of this bidirectional neural interface, in particular for monitoring infraslow and LFP activity, demonstrating that the recording performance is not compromised during stimulation.

### Active/passive hybrid device concept for graphene-based bidirectional neural interfaces

We developed a flexible epicortical device that features two intermixed arrays: nanoporous rGO stimulating microelectrodes and gSGFETs as active sensors, all fabricated at wafer scale (Fig. 1a- b, Fig. S1). In this device, the gSGFETs provide artefact-resilient, wide-bandwidth recordings, while the rGO microelectrodes offer high charge injection capacity for stimulation. The gSGFETs’ recordings can then be used to trigger or adjust the electrical stimulation delivered by the rGO microelectrodes, eventually enabling real-time, closed-loop neuromodulation (Fig. 1c).

**Fig. 1:**
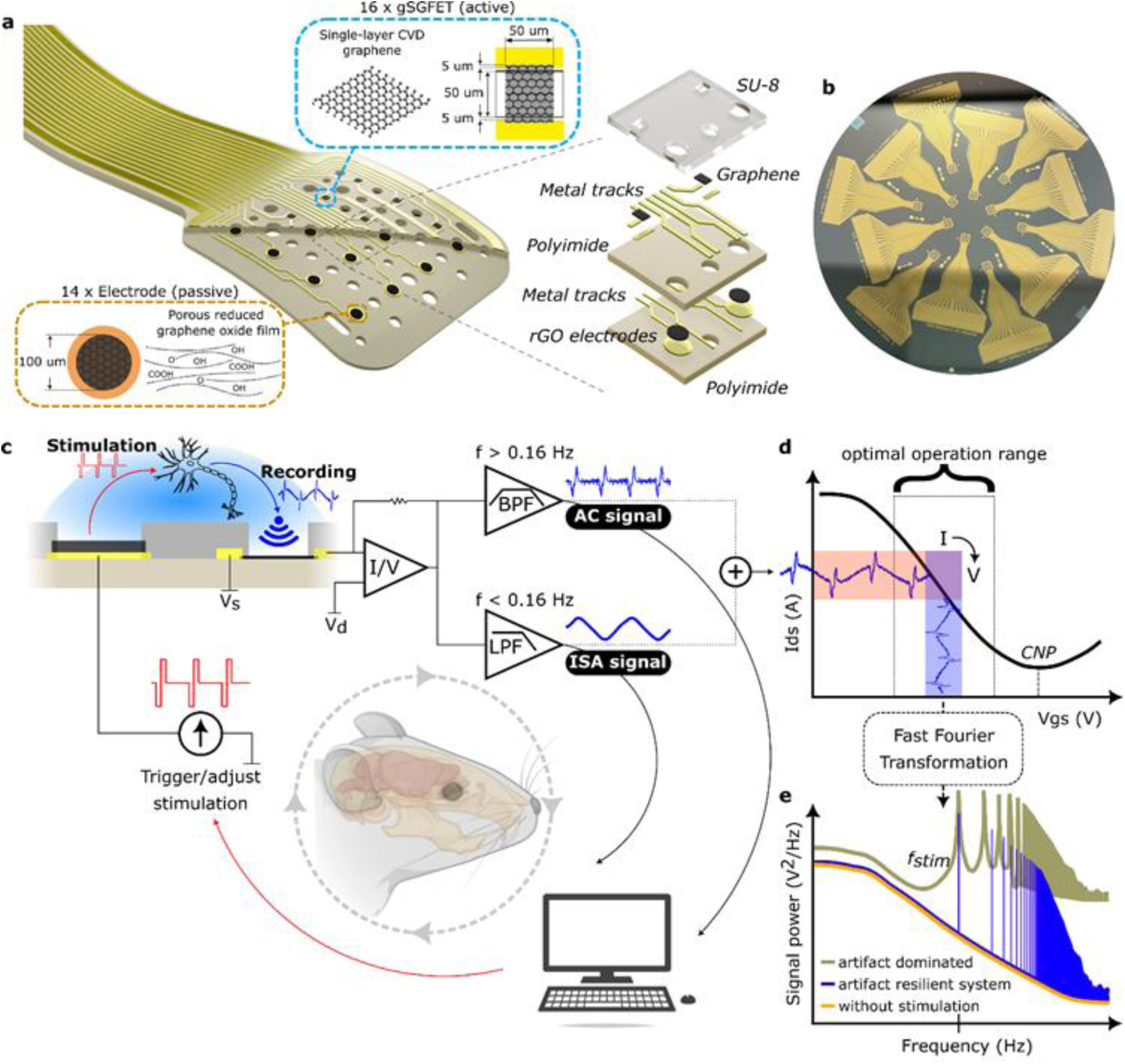
An artefact-resilient wide bandwidth bidirectional graphene neural interface. **a**, Tip of the flexible epicortical probe containing two intermixed arrays of porous reduced graphene oxide microelectrodes and graphene field-effect transistors. Polyimide provides insulation at the bottom and between the electrode and transistor levels while SU-8 covers the side of the probe facing tissue. **b**, Photograph of 4” wafer with 12 devices after fabrication. **c**, Closed-loop operation schematic of the active/passive hybrid device architecture. The microelectrodes enable electrical stimulation and the gSGFETs provide wide bandwidth recordings. The gSGFETs‘ ability to reliably record neural signals during simultaneous stimulation allows for precise triggering or adjustment of delivered stimulation in a closed-loop manner. **d**, Schematic representation of gSGFET current-to-voltage conversion. The gSGFETs’ ability to work in a wide gate voltage range, in addition to the tuneability of their operation point, allows them to be operated as active transducers in a wide dynamic range recording system. **e,** Power spectrum of signals acquired by artefact-dominated and artefact-resilient systems. In a recording system susceptible to stimulation artifacts, amplifier saturation can lead to contamination of the entire frequency spectrum.

A key technical challenge in neural implants is achieving simultaneous recording of neural activity while delivering stimulation. This is primarily due to the significant amplitude difference between neural signals and the stimulation-induced artefacts. Stimulation artefacts can reach amplitudes ranging from tens to hundreds of millivolts [35], whereas local field potentials (LFPs) are typically much smaller, often below 1 mV. Consequently, an amplification system with a large dynamic range is required to capture this range of signal amplitudes [34].

gSGFETs are well-suited for this task, as they can be operated as transducers with a very large dynamic range [36]. In a gSGFET (see Fig. 1c), the electrolyte in contact with the graphene channel functions as the gate of the transistor [27,28]. Variations in the electric potential due to neural activity couple to the channel as gate-to-source-voltage (Vgs) changes via the graphene- electrolyte interface capacitance, which alters the channel conductivity and results in drain-to-source current (Ids) changes. The Ids-Vgs curve represents the transfer function of the transducer (Fig. 1d). For gSGFETs, this transfer curve exhibits a V-shape due to the ambipolar transport in graphene, with the minimum of the curve corresponding to the charge neutrality point (CNP). Away from the CNP, the current is eventually limited by the metal-graphene contact resistance [36]. The transfer curve defines the transistor’s dynamic range (Fig. 1d), typically limited to the linear part of the curve, which can be optimized by tuning the Vgs applied bias.

The transduced signal is then amplified, as described in prior works [20,36,37]. Briefly, the system consists of two stages (Fig. 1c): in the first stage, a transimpedance amplifier converts the current signal into a voltage signal. The second stage further amplifies the voltage signal before digitalization. To ensure proper signal acquisition across the wide range of signal amplitudes (from low-amplitude LFPs to large infraslow potential variations and drifts), the signal is divided into two frequency bands: the AC signal and the ISA signal (Fig. 1c) with high and low gains, respectively. The ISA signal is defined by a low-pass filter (LPF, < 0.16 Hz) while the AC signal is defined by a band-pass filter (BPF, 0.16 - 10 kHz). The total signal is then reconstructed in post-processing by summing the signals from the two bands and calibrating them through interpolation into the transistor’s transfer curve, yielding distortion-free DC-coupled recordings [36].

While the above-described amplification strategy based on signal decomposition into two frequency bands is not exclusive of transistor-based systems, transistors offer several advantages that contribute to the resilience of this technology to stimulation-induced artefacts. First, the current limitation of the transistor away from the linear regime protects the transimpedance amplifier from saturation, even if the recorded signal exceeds the transistor’s dynamic range. Additionally, the dynamic range of the second-stage amplifier can be fixed to match the expected range of drain-source current values, thus avoiding saturation. Second, gSGFETs can be repolarized in response to large drifts, optimizing their dynamic range. In addition, the elimination of the high-pass filter commonly employed with passive electrodes, which is necessary to enable DC-coupled recordings, facilitates rapid recovery in the unlikely event of saturation [34]. These features collectively form the basis of the high-fidelity signal recording during stimulation of this system (Fig. 1e), as demonstrated experimentally in the results presented below.

### Device fabrication and characterization

The flexible epicortical design used in this study consists of two intermixed arrays of rGO microelectrodes (circular, 100 μm diameter) and gSGFETs (squared channel, 50x50 μm^2^) (Fig. 2a,b). Both arrays have 4 rows and 4 columns (450 μm pitch) and are shifted by 225 μm in relation to each other. The entire array covers a total area of 1.5 mm x 1.5 mm. The fabrication begins with a sacrificial 4’’ SiO_2_/Si wafer substrate on which a thin layer of polyimide (PI) is deposited (Fig. 2b). Subsequently, the rGO microelectrode layer is fabricated as previously reported [29,30,31]. A second PI layer encapsulates the microelectrode layer providing electrical insulation with the transistor layer. After definition of the VIAs (vertical interconnect accesses) to electrically connect the microelectrode layer with the transistor layer, the gSGFETs are fabricated and passivated with SU-8 resist which acts as device’s top encapsulation. Finally, the second PI layer is etched to expose the microelectrodes underneath. A detailed description of the fabrication protocol is provided in the Methods section and is schematically represented in Fig. 2b.

**Fig. 2:**
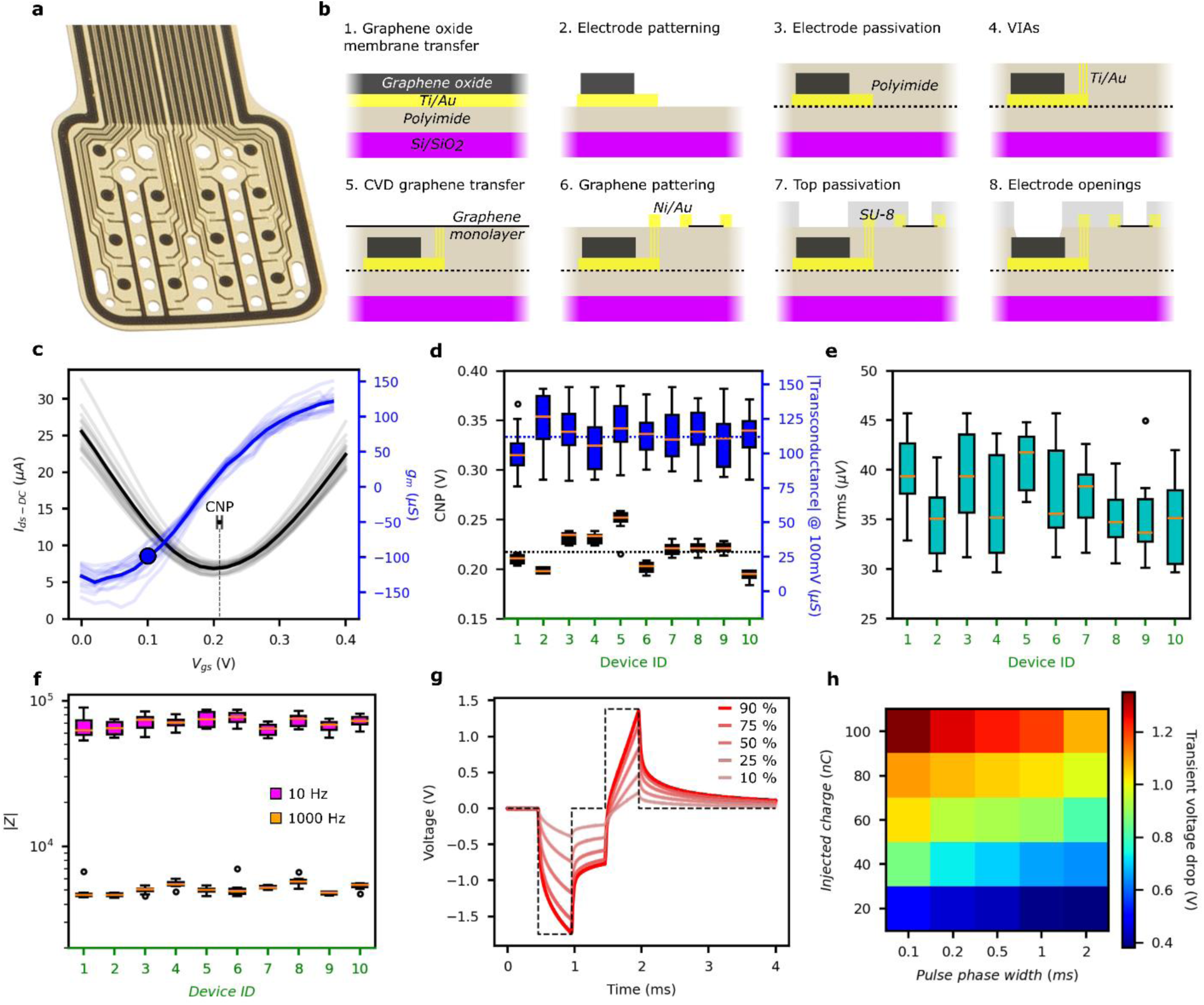
Characterization of hybrid arrays. **a**, Photograph of a tip of a flexible epicortical probe. **b**, Microfabrication steps for the monolithic integration of the transistor and microelectrode technologies. **c**, Individual transfer curves (faint black lines) and transconductance (faint blue lines) of 16 graphene transistors in one device. Thicker lines correspond to the average of the individual transistor curves. **d,** CNP and transconductance values (for Vgs = CNP – 0.1V) dispersion statistics for 10 devices (16 transistors in each). **e**, Transistor noise dispersion statistics for 10 devices. **f**, Impedance values at 10 and 1000 Hz for 10 devices (14 electrodes in each). **g**, Voltage response measured by a stimulating rGO microelectrode during application of current-controlled biphasic pulses of 500 μs (dashed lined) with increasing current amplitude; the numbers in the inset correspond to a percentage of the maximum current amplitude before the electrode is operated outside the potential window. **h**, Cathodic capacitive voltage shift measured by a rGO microelectrode during the injection of current pulses at different levels of injected charge and pulse widths.

Verification of technology integration and device quality tests were conducted in standard 1x phosphate-buffered saline (PBS) solution. The electrical characterization of the gSGFETs (see Fig. 2c-e) reveals that the performance of the gSGFETs integrated in the hybrid devices is within the expected range in terms of transconductance and CNP when compared with previously published data from non-hybrid devices [21,22,23]. The CNP values are homogenous within the same device and usually are only slightly offset between different microfabrication batches (Fig. 2d), which we tentatively attribute to uncontrolled doping during graphene transfer and device processing. For 160 measured transistors from various batches, the average CNP value is 0.2182 V with a standard deviation (**σ**) of 0.017 V. Similarly, the transconductance (dIds/dVgs) is also homogenous across multiple batches with a mean of 113.26 μS (**σ** <19.09 μS) for all transistors. The transistors’ sensitivity has been assessed using the effective gate noise (Vrms) calculated as the root-mean-square of gate voltage noise integrated between 1Hz and 2kHz, with average values of 33 to 43 μV (Fig. 2e).

The performance of microelectrodes is also homogenous across multiple devices. Electrochemical impedance characterization reveals an average impedance of 5.1 kΩ (**σ<** 0.5kΩ) at 1kHz (Fig. 2f and Fig. S2d). The maximum current that could be safely injected with the rGO microelectrodes was established by finding the biphasic pulse amplitude that elicited an electrode interfacial voltage that remains within the limits of rGO’s electrochemical water window (−0.9 V for cathodic and +0.8 V for anodic versus Ag/AgCl reference) (Fig. S2e). As discussed in Supplementary Information (Fig. S3), the ohmic drop is used to calculate the interfacial voltage. Considering the maximum safe pulsed current amplitude (which we refer to as “Amax”), we characterized the voltage polarization at the electrode interface in response to biphasic pulses of varying pulse width (0.1 - 2 ms) and for a range of amplitudes (referred to 10 - 90% of Amax, typically in the range 20-390 μA, see Fig. S3c) (Fig. 2g-h). All in all, the characterization results show that the developed monolithic integration microfabrication process maintains the performance of gSGFETs and rGO microelectrodes, comparable to their performance when fabricated separately [21,22,23,33].

### gSGFET recording-performance characterization during stimulation

#### In vitro assessment of gSGFET recording performance during stimulation

The recording capabilities of gSGFETs, on the one hand, and the recording and stimulation capabilities of rGO microelectrodes, on the other hand, have been described separately in previous publications [20,21,22,23,26,27,28,31,32,33]. In this study, we evaluated the performance of monolithically integrated hybrid devices, with a focus on their operation involving simultaneous stimulation and recording with the same neural probe.

Stimulation artefacts typically consist of large voltage transients resulting from the current injected through the stimulation electrode [34]. The injected current spreads through the conductive medium (tissue or electrolyte), generating a transient electric potential whose amplitude is position-dependent: higher near the stimulation electrode and decreasing away from it. When the artefact amplitude is large, it can saturate the signal acquisition chain, compromising the quality of the recorded signal.To assess the signal acquisition capability of the hybrid devices during stimulation, we designed a protocol to test a broad range of stimulation parameters, including pulse width (100 or 500 μs per phase), stimulation frequency (25, 130, 185, 250 Hz), and current amplitude (10, 25, 50, 75, 90 % of Amax) (Fig. 3a). The stimulation protocol was applied to one electrode in the array, and the effect of the stimulation on the quality of recordings was assessed across the entire array (both for transistors and electrodes).

**Fig. 3:**
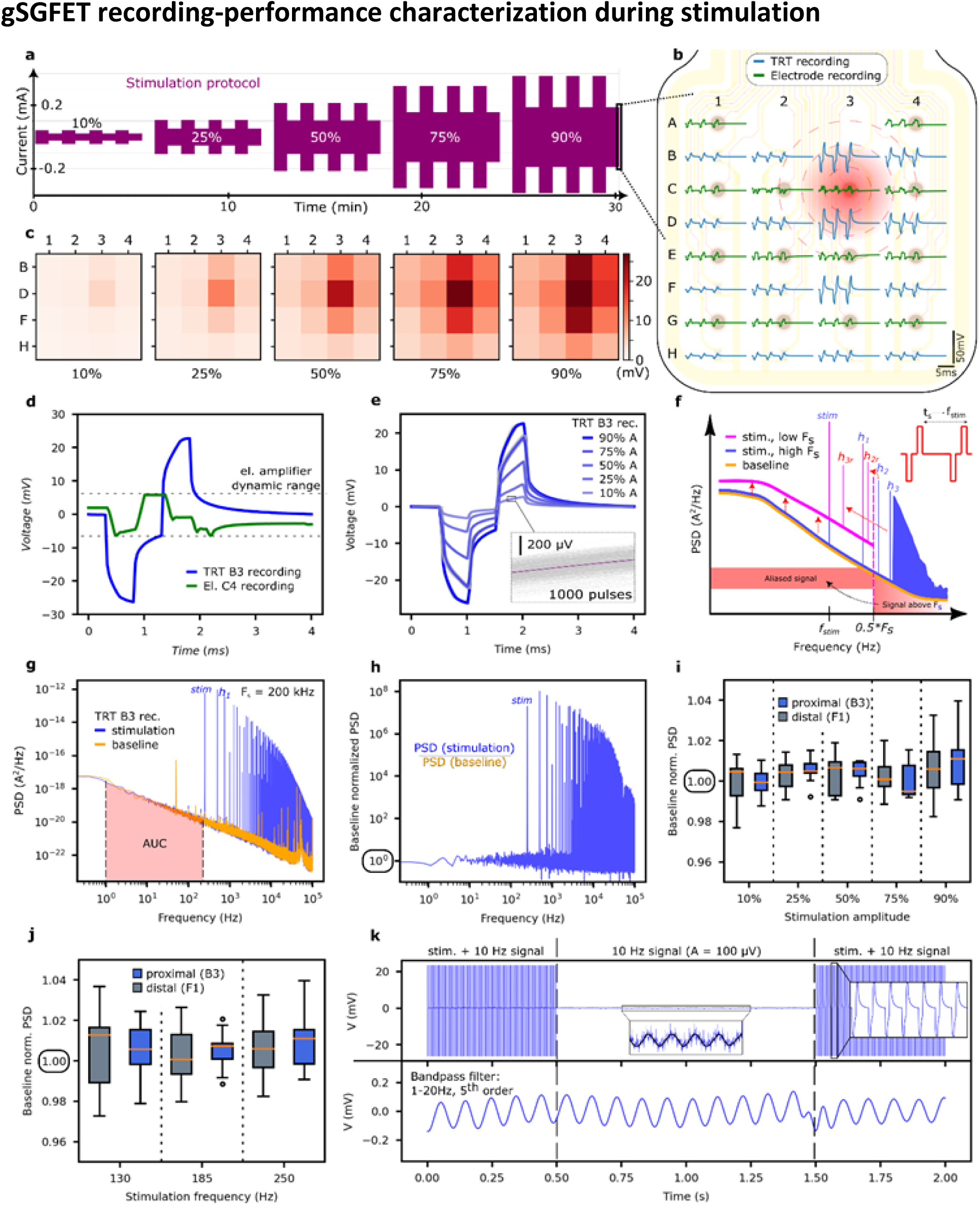
In vitro assessment of recording performance of gSGFETs during stimulation. **a**, Stimulation protocol with combinations of various stimulation parameters: pulse width (100 or 500 μs per phase), frequency (25, 130, 185 or 250 Hz), amplitude (10, 25, 50, 75 or 90 % Amax). Each configuration lasts ∼40 s. **b**, Electrode (green line) and transistor (blue lines) recordings. Mapping of the last 3 pulses (250 Hz, 500 μs/phase, 90% Amax) in the stimulation protocol. Stimulation through electrode C3. **c**, Stimulation induced voltage mapped by transistors. **d**, Comparison of the 500 μs stimulation-artefact pulse recorded by a transistor and an electrode showing electrode amplifier saturation **e**, Transistor recordings of 1000 stimulation-artefact pulses (F=250 Hz, t=500 μs, A=90%); the inset shows the 1000 pulses overlapped. **f**, Schematic of noise aliasing and signal quality dependence on sampling frequency. Stimulation causes high frequency harmonics which, due to aliasing, contaminate the frequency band of interest. Using high sampling frequency reduces the aliasing effect, as shown by the PSD representation. **g**, Comparison of the PSD for the recording of a transistor in PBS with (blue) and without (orange) stimulation (F=250 Hz, t=500 μs, A=90%). **h**, Ratio of the PSDs from panel g, defined as baseline-normalized PSD. **i**, Statistics of baseline-normalized PSD value as a function of the stimulation amplitude. For each amplitude, the data corresponds to the average of 9 full stimulation protocols (as shown in panel a). **j**, Same as in panel i, but as a function of the stimulation frequency. **k**, Raw recording and recovery of a 10 Hz signal during stimulation.

Fig. 3b illustrates a map of the array with representative voltage traces recorded with transistors and electrodes for a particular stimulation protocol applied to one of the electrodes (C3 in Fig. 3b). Thanks to the large dynamic range of gSGFETs, these devices can record the voltage change induced by the stimulation current pulse. As shown in Fig. 3d (see also Fig. S4 and S5 in Supplementary Information), the gSGFETs accurately measure the stimulation-induced potential change. Fig. 3e depicts voltage traces recorded by a transistor next to a stimulating electrode at various stimulation amplitudes. In this plot, the response to 1000 consecutive pulses is overlapped, confirming that each stimulation pulse produces the same response as measured by the gSGFETs, in contrast to the case of the electrodes, which are not able to record the stimulation-artefacts faithfully. It is worth noting that the signal measured by the transistors consists of two components, an ohmic drop component arising from the injection of biphasic current pulses into a conductive medium, and a component that resembles the charging of a capacitor by the biphasic pulse. We tentatively attribute this second component to pA-level leakage currents between the stimulating and the recording electronic systems. Under the tested stimulation conditions, the recorded signal amplitude by transistors a few hundred micrometers from the stimulating electrode reaches tens of millivolts. As expected, the signal amplitude decreases with increasing distance from the stimulating electrode, as shown in Fig. 3c.

Once we have validated the fidelity of the transistors in recording the stimulation-induced pulses, we will evaluate the level of perturbation introduced by the stimulation in the recorded signals, by performing a frequency domain analysis. Fig. 3g shows the power spectral density (PSD) of signals measured by one of the transistors with and without stimulation, using a sampling frequency of 200 kHz. Figure 3h depicts the ratio of the PSDs before and during stimulation, representing a baseline-normalized power spectral density. The PSD of the baseline signal without stimulation displays the characteristic 1/f noise of gSGFETs. When stimulation is applied with a neighboring microelectrode, the PSD of the recorded signal shows a peak at the fundamental stimulation frequency (f_stim_ = 130 Hz) and its corresponding harmonics. Even for the gSGFETs, high-frequency harmonics associated with the stimulation artefact, combined with the limited sampling frequency, can lead to signal contamination through aliasing. As illustrated in Figure 3f, aliasing mitigation requires careful adjustment of the antialiasing filters and selection of the sampling frequency in relation to the fundamental frequency of the stimulation pattern. In order to quantify the level of contamination introduced by the stimulation-induced artefact in the frequency range below the stimulation frequency, we calculated the root mean square (RMS) of the recorded signal in the 1-125 Hz frequency range and computed the ratio of the RMS before and during stimulation, which could be used as indication of distortion. Fig. 3i and Fig. 3j summarize the RMS ratio as a function of stimulation amplitude and stimulation frequency, respectively. Data were calculated based on 9 separate runs of the full stimulation protocol shown in Fig. 3a. For all tested stimulation parameters, the RMS ratio was 1 ± 0.04, indicating no significant difference between recording with and without stimulation below the stimulation frequency. Additionally, we verified that there was no difference in signal quality between recordings made by transistors that were proximal (closest to the stimulating electrode) or distal (furthest from the stimulating electrode), as shown in Fig. 3i and Fig. 3j.

As demonstrated above, in the case of the gSGFETs’ recordings, the frequency range below the stimulation frequency is not altered by the artefact, which allows the use of simple frequency filtering to remove the contribution of the stimulation-induced artefact. Fig. 3k demonstrates the recovery of an externally applied test signal during stimulation using just signal filtering of the raw data. In this experiment, a 10 Hz sinusoidal wave with 100 μV amplitude, simulating an LFP signal, was applied to the saline solution in which the hybrid array was tested. The top panel in Fig. 3k shows the raw data (without filtering) of the recording from one transistor during and after stimulation, while the bottom panel shows the same recording after applying a digital bandpass filter (1-20 Hz, 5^th^ order). This experiment demonstrates that even during stimulation, when large-amplitude stimulation artefacts are present in the recordings, the gSGFETs can recover the test signal with high fidelity.

#### In vivo assessment of gSGFETs recording performance during stimulation

To assess the artefact resilience of the device *in vivo*, acute experiments were performed in awake head-fixed mice (n=4) (see Fig. 4a and Methods). The device was placed over the right somatosensory cortex with electrode and transistor references in the contralateral hemisphere (see Fig. 4b). The experimental paradigm consisted of a series of stimulations, each of 20 to 30 seconds of duration. 100 Hz, 21 μA (10% of Amax), 500 μs/phase biphasic pulses were applied (Fig. 4c). In order to demonstrate the ability of the gSGFETs to capture relevant biological signals we injected a chemoconvulsant (picotroxin, 10 mM) prior to electrical stimulation to induce large amplitude epileptiform activity (Fig. 4e). Following picotroxin injection, epileptic discharges developed as shown by the time series and in the frequency spectrum of an example transistor channel (see Fig. 4e). During subsequent stimulation, see Fig. 4f, the frequency component of the epileptiform activity remained unaffected as recorded by the gSGFET, showing the robustness of gSGFETs recordings to in vivo electrical stimulation artefacts. The time domain representation of the period around the stimulation (see Fig. 4g) also reveals that, after low-pass filtering (<60 Hz), the recording of the epileptic activity was preserved during stimulation, in proximal as well as distal channels (colored maps in Fig. 4g).

**Fig. 4:**
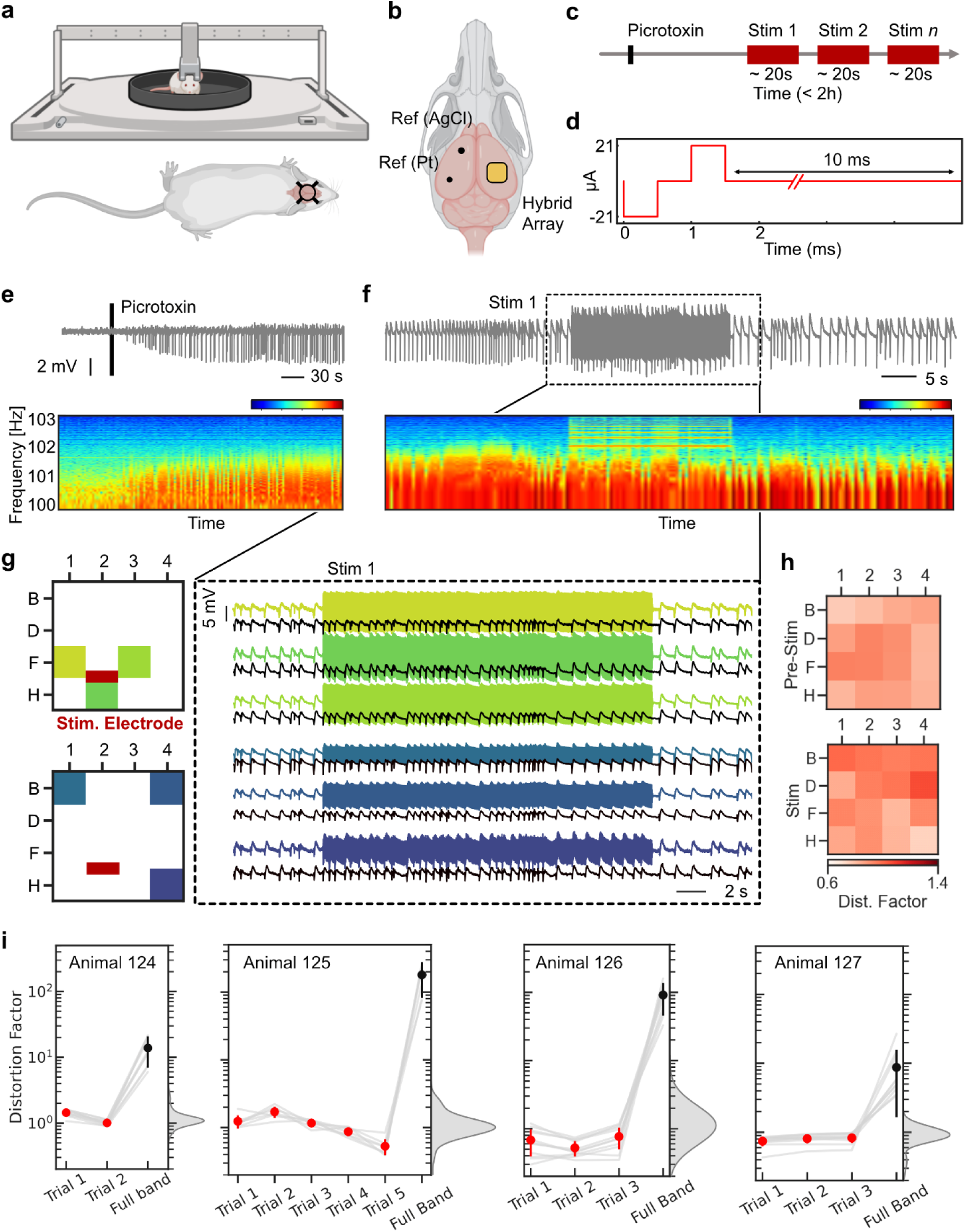
***In vivo* gSGFETs recording performance during electrical microstimulation**. **a**, Schematic drawing of mouse on Neurotar head-fixation system with a blow up (**b**) of the top cranial view showing the hybrid array position relative to ground and reference wires. **c**, Experimental timeline with indication of the picrotoxin chemo convulsant injection and subsequent stimulations. **d,** Detail of biphasic stimulation protocol. **e-f**, Voltage recording (grey) and time-frequency analysis of an example transistor channel before and after picrotoxin injection (e) and before after electrical stimulation (f). Note the limited spread of stimulation bands leaving biological activity of lower frequency unaffected. **g**, Detail of the first stimulation period across 3 proximal and 3 distal channels with full-band (colored traces) and low-pass filtered at 60 Hz activity (black traces). **h**, Spatial colormap of the RMS ratio, represented as a distortion factor, before and during stimulation for all transistor channels. **i**, RMS ratio across consecutive trials (red) of 4 animals with the averaged full band values (black) and kernel density estimate of the surrogate values in the absence of stimulation.

To validate these results and determine the degree to which the signal is affected by stimulation, we quantified the artefact induced on the LFP band (<60Hz) employing the previously described RMS ratio (see also Fig. S6 for an *in vivo* example of the RMS ratio), which provides information about how the stimulation distorts the recorded biological signal. Fig. 4h presents the distortion factor immediately before and during stimulation across working transistors using the mapping presented in Fig. 3. Note that, although RMS ratio values are similar before and during stimulation, as well as clustered around one (i.e. no distortion), there is considerably more variance compared to the in vitro experiment (see Fig. 3i,j). We hypothesized that this may be due to inherent variability of the biological signal. Therefore, to assess the distortion factor across all animals while accounting for the physiological changes in the biological signal, we computed the RMS ratio for each stimulation trial as well as a surrogate distribution using consecutive periods without stimulation. We then present this distribution as a kernel density estimation (KDE) which represents the variability of distortion factor values in the absence of stimulation. As shown in Fig. 4i, RMS ratio values below 60 Hz during stimulation trials fall within the limits of the surrogate distribution (gray KDE plot) and are considerably lower than the average full band RMS ratio of each trial. Altogether this demonstrates that, below the stimulation frequency, the gSGFETs present a negligible degree of distortion over baseline activity across trials and animals, enabling high-fidelity recording of the underlying relevant biological activity even during stimulation.

#### Proof-of-concept applications of the hybrid neural interface

Having assessed the stimulation-induced artefact resilience of the device both in vitro and in vivo, we designed a series of experiments to demonstrate the advantages of hybrid arrays in the context of recording infraslow activity during stimulation and infraslow-based neuromodulation. Cortical spreading depolarizations (CSDs) are slowly propagating waves of neuronal and glial depolarization associated with a number of neurological disorders including epilepsy, stroke and migraines with aura [38, 39]. Here, we demonstrate the ability of the hybrid neural interfaces to induce and record CSDs with high spatiotemporal resolution. Fig. 5a depicts the timing of two stimulation trains applied with the rGO microelectrodes (20 Hz, 21 μA biphasic pulses, 5 seconds), and the corresponding induced CSDs measured with the gSGFETs. These electrically induced SDs present the expected silencing of higher frequency activity (see Fig. S7a) and exhibit homogeneous values of amplitude and duration across animals (see Fig. S7b). A detailed view of the first event is presented in Fig. 5b, which shows the differences in CSD onset between transistors. All channels are sorted, and color coded depending on the CSD arrival at each transistor. The inset in Figure 5b shows the spatial layout of these channels and demonstrates how transistors closer to the stimulation site (red rectangle in Figure 5b) detect the CSD earlier than those far from it. A more detailed view of the relationship between detection onset and distance of transistor to stimulation electrode is presented in Fig. 5c; the correlation values of the linear regression between channel distance to stimulation site and CSD onset are r=0.82, and p=1.9e-4. Finally, Fig. 5d shows how this relationship is consistent within and across animals, proving that the hybrid array can be used to reliably induce and detect infraslow brain dynamics.

**Fig. 5.**
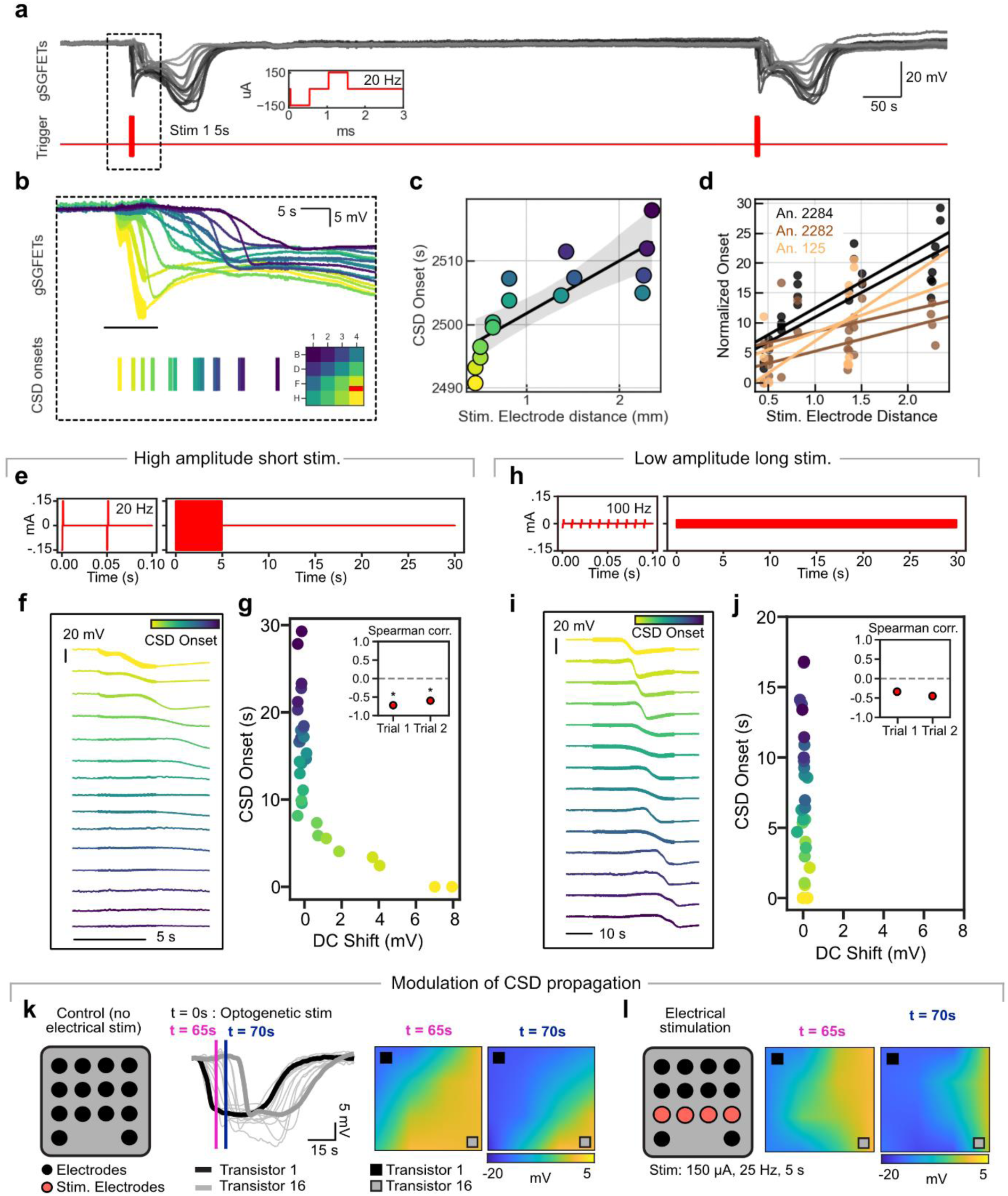
Proof-of-concept applications of the hybrid neural interface. **a**, Reliable induction of CSDs by electrical stimulation with rGO microelectrodes. Inset shows stimulation train protocol of biphasic pulses at 20 Hz for 5 seconds. **b**, Detail of the initiation of the CSD (upper panel) with color coded channels matching the beginning of the CSD. Note on the inset that the CSD originates on the lower right corner of the device, near the stimulation site (red). **c**, Linear regression between the CSD onset of panel b and the channel distance to stimulation site. **d**, Linear regressions across trials and animals. **e-g**, DC shifts induced by high amplitude short stimulation. **e**, Stimulation protocol. **f**, DC shifts observed during electrical stimulation are only present in channels close to CSD onset. **g**, Relationship between CSD onset and DC shifts reveals a significant correlation (Spearman rank test) for two stimulation trials (*p < 0.05). **h-j**, DC shifts during low amplitude long stimulation. In this scenario, responses do not present DC shifts and thus no relationship is present between CSD onset and DC shift. **k**, Control spread of an optogenetically-induced CSD in the absence of electrical stimulation. **l**, Modulation of the direction of CSDs propagation due to the electrical stimulation from a row of electrodes.

Given the ability of the hybrid array to simultaneously record infraslow activity in the presence of electrical stimulation, we aimed to study if this technology would allow us to infer information about the underlying mechanism through which CSDs are induced. We employed two stimulation paradigms, one of high amplitude and short duration (Fig. 5e-g) and one of low amplitude and long duration (Fig. 5h-j). Only in the first paradigm, we observed DC shifts near the stimulating electrode that preceded the onset of CSD. This underscores a key advantage of the technology, especially for studying CSDs associated with seizures. As neurons fire synchronously during a seizure, they release large amounts of extracellular potassium, which is reflected in the DC shift. Unlike LFP signals, which can spread over a broader area, the DC shift provides more localized information about the seizure’s origin. This insight is only possible due to the hybrid array’s ability to record full-band activity during periods of electrical stimulation.

The potential to study CSDs may also hold translational promise provided their involvement in a number of neurological conditions. Therefore, the ability to modulate its propagation through cortical tissue may be of use for future therapeutic avenues. As a first approach to demonstrate the capability of our hybrid technology to detect and modulate CSD propagation, we used a previously described *in vivo* model where CSDs could be induced on demand optogenetically [22] (see Methods and Fig. 5e-f). Figure 5k shows an example of the propagation of a CSD in the absence of electrical stimulation. Then, to assess if its propagation direction could be modulated, we repeated the optogenetic stimulation but applied electrical stimulation (150 uA, 25 Hz, 5s from 4 electrodes) right after CSD onset. We present two timeframes (t=65 s and t=70 s) from the onset of optogenetic stimulation (t=0 s). Notice the linear (radial)trajectory in Fig. 5k, something typically seen in optogenetically induced SDs [22] and the comparison with Fig. 5l, where the propagation seems to be facilitated in the direction of stimulation.

As an additional translational application, we assessed the device’s performance at mapping out epileptogenic tissue. For a subset of patients with drug-refractory epilepsy, surgical resection of the epileptogenic tissue is a therapeutic option but only when it can be accurately identified prior to surgery. One way to identify epileptogenic tissue is by monitoring the tissue response to neurostimulation [40]. Stimulation that results in time-locked responses in healthy tissue, can result in after stimulation discharges, seizures, or SDs when applied to hyperexcitable epileptic tissue [41]. Therefore, to model a potential future clinical scenario for this technology we performed experiments in a mouse model of neocortical epilepsy based on conditional knock- out of the *Tsc1* protein unilaterally in the somatosensory cortex (see Fig. S8a-b). The model led to spontaneous seizures and hyperexcitable areas. (see Fig. S5c). During the experiment, the hybrid array was positioned to cover areas within and outside the hyperexcitable site. We then stimulated on and off this area and observed differential responses in the form of post-stimulation epileptiform discharges and SDs that only occurred when stimulating on the targeted area (see Fig. S8d). To the best of our knowledge, this is the first demonstration of simultaneous focal stimulation and ISA recordings in the context of mapping epileptogenic tissue.

Altogether, the hybrid array’s ability to induce and modulate infraslow activity with spatiotemporal accuracy could become a key feature of this technology in the investigation, diagnosis, or treatment of a wide range of neurological disorders.

## Conclusions

In this work, we demonstrated the monolithic integration of passive nanoporous graphene electrodes and graphene field effect transistors in a single device. The technology was developed on 4” wafers and can be easily scaled up to larger wafer sizes without significant changes. This hybrid neural interface combines the advantages of passive electrodes and transistors while maintaining their performance, i.e. high density microstimulation, and wide-bandwidth recordings. We validated the technology in vitro by testing a wide range of stimulation parameters, showing that gSGFETs recordings are resilient to stimulation-induced charging allowing wide-band recordings during stimulation. Moreover, we investigated several proof-of-concept applications in vivo, proving the device’s ability to induce and modulate infraslow activity with spatiotemporal accuracy as a promising feature for the investigation of a wide range of neurological disorders. In epilepsy, ISA and DC-coupled recordings could allow more comprehensive assessment of the pathophysiological processes associated with either an increase in neuronal activity during seizures or a loss of neuronal activity during a spreading depolarization (SD). The ability to record ISA together with higher-frequency activity would facilitate the understanding and treatment of pathological states. The integration of information acquired from the entire spectrum of brain electrophysiological signals could enable new applications and enhance outcomes of existing therapies by defining novel biomarkers based on a full-band characterization of the pathological states.

Overall, this work describes a universal tool for neuroscientific research with high-current-density microsimulation and artefact-resilient full bandwidth recording capabilities that could be used to investigate the functional role of wide frequency band brain activity with unprecedented spatiotemporal resolution. The proof-of-concept applications (spreading depolarization assessment and neuromodulation, and wide-band epileptogenic mapping) show the potential of our technology for clinical translation. We envision that further development of this technology will lead to infraslow biomarkers being used in closed-loop systems, alone or in combination with higher frequency biomarkers, leading to improved efficacy of adaptive patient-specific neuromodulation.

## Methods

### Graphene growth

Graphene monolayers were obtained by chemical vapor deposition growth on a 4x8 cm^2^ copper foils (foil thickness – 0.035 mm). The foils were electropolished in a solution containing H_2_O (1 l), H_3_PO_4_ (0.5 l), ethanol (0.5 l), isopropanol (0.1 l) and urea (10 g) prior to the graphene growth. After polishing, the foils were placed in a quartz tube (1600 x 60 mm^2^) and heated in a three-zone oven. The first annealing step at 1050 °C under a 400 sccm argon flow at 100 mbar for 1.5 h was followed by a 20 min growth step at 12 mbar under a gas mixture of 1000 sccm argon, 200 sccm hydrogen and 2 sccm of methane. The sample was then cooled under a 400 sccm argon flow by removing the quartz tube from the oven.

### Graphene characterization

All graphene/Cu sheets were characterized with Raman (Witec spectrograph). 30x30 µm^2^ maps were obtained with a sub-um resolution and 488 nm laser excitation wavelength to minimize copper substrate luminescence signal. The laser power was set to 1.5 mW. A 600 g/mm grating was used to provide a pixel-to-pixel spectral resolution below 3 cm^-1^.

### GO membrane preparation

Aqueous GO solution (Angstron’s Materials) was diluted in deionized water to obtain a 0.15 mg/ml solution and vacuum filtered through a nitrocellulose membrane with pores of 0.025 µm, forming a thin film of GO [31]. The thin film was then transferred to the target substrate using wet transfer in deionized water and further thermal annealing at 100 °C for 2 min. The GO film–substrate stack was hydrothermally reduced at 134 °C in a standard autoclave for 3 h.

### Fabrication of hybrid arrays

Hybrid arrays were fabricated using microelectronic fabrication techniques at the IMB-CNM Clean Room on sacrificial 4” Si/SiO_2_ wafers. First, the wafers were cleaned in Piranha bath to remove organic contamination present on the surface. Oxygen plasma was applied to clean the surface and increase adhesion to polyimide (PI-2611, HD MicroSystems) which was then spun at 1500 rpm to form a 7,5 µm-thick layer after 350 °C bake in nitrogen atmosphere. 20 nm of Ti and 200 nm of Au were sputtered on the PI surface. Oxygen plasma was applied before GO membrane transfer, which was prepared as described above. A photolithography with an AZ nLOF 2070 (MicroChemicals) photoresist was performed to define microelectrodes followed by e-beam evaporation of 80 nm of Al and lift-off process. After the lift-off, rGO film was etched with oxygen reactive ion etching everywhere except the areas where it was protected by Al. The Al layer was then removed with standard aluminum etchant (phosphoric and nitric acid solution) revealing 100 µm diameter electrode discs. A photolithography was performed to define shapes of electrode tracks in Ti/Au metal layer. The wafers were first immersed in a commercial gold etchant solution (TechniEtch ACI2, Microchemichals) for 2 min and a mixture of propandiol:H2O:HF:50% for 1 min to etch titanium in areas not protected by the photoresist. Low power oxygen plasma was used to clean wafer surface and increase adhesion of second polyimide layer which was spun at 5000 rpm to form 2,5 µm-thick insulating film over the electrode layer. After a 350 °C bake of the PI in nitrogen atmosphere, Vertical Interconnect Accesses (VIAs) were patterned by photolithography with AZ 10xT positive resist (MicroChemicals). Oxygen reactive ion etching was used to etch through polyimide and expose electrode tracks underneath. The AZ 10xT residues were removed and photolithography with image reversal AZ 5214 resist (MicroChemicals) was done before evaporation of 10 nm of Ti and 100 nm of Au to form VIAs and first metal layer for graphene transistors located on top of the second polyimide layer over rGO electrode level. After a lift-off process, the wafer surface was treated with low power oxygen plasma to enhance adhesion with single-layer graphene (SLG) which was transferred right after using wet transfer method. Subsequently, the SLG was patterned with positive photoresist photolithography (HiPR 6512, FujiFilm) and RIE (1 min, oxygen plasma). After wafer cleaning (20 min in acetone -> 10 min in isopropanol -> 5min in DI water) another photolithography (image reversal AZ 5214 photoresist, MicroChemicals) was performed to define second metal layer for graphene transistors. Prior to evaporation of 20 nm of Ni and 200 nm of Au, 20 min of UVO treatment was applied to improve graphene/metal contacts. After the lift-off, SU-8 2005 photoresist (Kayaku Advanced Materials) was spun on the wafer at 3000 pm (2 µm-thick layer), exposed, developed and hard baked at 120 °C for 20 min. Top SU-8 passivation was followed by AZ 10xT (MicroChemicals) photolithography which defined the electrode openings. Polyimide over rGO electrodes was etched with RIE in order to create electrode openings. After wafer cleaning in isopropanol (10 min), ethanol (30 min) and DI water (5 min), the last photolithography with AZ 10xT photoresist to define device outlines and holes was done. After RIE of the two polyimide layers, the wafer was cleaned, and the flexible devices manually peeled off from the sacrificial Si/SiO_2_ wafer.

### Characterization of the hybrid arrays – rGO electrodes

Electrochemical characterization was performed using a potentiostat (Metrohm Autolab PGSTAT128N) in a three-electrode configuration. An Ag/AgCl electrode (FlexRef, WPI) was used as the reference electrode, and a platinum wire (Alfa Aesar, 45093) as the counter-electrode. The devices were immersed in a solution prepared by dissolving one PBS tablet (Sigma-Aldrich, P4417) in 200 ml of distilled water. The final solution contains 10 mM phosphate buffer, 137 mM NaCl, and 2.7 mM KCl at pH 7.4. Before the in vitro electrochemical evaluation, the electrodes underwent an electrochemical activation process consisting of 33 cyclic voltammetry cycles in the –0.9 V - 0.8 V range at 50 mV/s rate.

Electrochemical characterization in PBS involved measuring impedance, cyclic voltammetry, and current pulses to evaluate functionality, ensuring no broken or electrically shorted electrodes. Electrode impedance spectroscopy was conducted using the previously mentioned three-electrode setup by applying a 10 mV sinusoidal wave over a frequency range from 1 Hz to 100 kHz.

### Characterization of the hybrid arrays – gSGFETs

The gSGFETS were characterized together with the electrodes in the same 150 mM PBS solution. Drain–source currents (*I*_ds_) were measured varying the gate–source voltage (*V*_gs_) versus a Ag/AgCl reference electrode, which was set to ground. Steady state was ensured by acquiring the current only after its time derivative was below a threshold (10^−7^ A s^−1^). The detection limit of the gSGFET was assessed by measuring the PSD of the DC current at each polarization point *V*_GS_. Integrating the PSD over the frequencies of interest (1 Hz–2 kHz) and using the transconductance allowed us to calculate *V*_RMS_.

### Animals

Different strains of mice were employed according to the Animal Act 1986 (United Kingdom Scientific Procedures) and under the licenses PPL PAF2788F5-13691 and PIL code 18068, endorsed by the Home Office. Animals were housed collectively as litter mates in 12 hour light/dark cycles prior to injection and head-bar surgery, after which housing was done individually. Water and food were provided throughout the day ad libitum. In vivo device performance experiments and CSD modulation experiments were carried out on wildtype (WT) C57BL/6J mice (4-6 months) while epileptogenic mapping experiments were performed using conditional Tsc1-floxed mice (Jackson Laboratory) cross-bred to obtain homozygotic specimens.

To obtain a focal knockout of Tsc1 in astrocytes, animals were injected with a viral construct (AAV5) containing a Cre recombinase and a glial fibrillary acidic protein (GFAP) promoter. Additionally, a fluorescent tag (mCherry) was added to confirm expression (AAV5-GFAP105- mCherry-Cre, AddGene, USA). CSD modulation experiments were carried out on C57BL/6J mice injected with a viral vector (AAV9) containing ChR2 and an enhanced yellow fluorescent protein (EYFP) tag to confirm expression once again (AAV9-CamK2a-ChR2-EYFP, AddGene, USA). Selective ChR2 expression in excitatory glutamatergic neurons was achieved through a Calcium/Calmodulin Dependent Protein Kinase II Alpha (CamK2a) promoter.

### Surgeries

Sterile environments and aseptic tools were common to all surgeries. Anesthesia was delivered through an isoflurane enriched chamber (1-1.5%; Henry Schein) prior to placing the animals on a stereotaxic frame (Kopf Instrumentation, USA). At the beginning of each procedure, pain relief medication was administered subcutaneously in the combined form of Meloxicam (15 mg/kg), and Buprenorphine (0.5 mg/kg). At the closure of the surgery, warmed subcutaneous saline (0.015 ml/g) was applied at body temperature to avoid thermal shock and provide hydration. For head-plate positioning, a single hole was drilled into the skull in proximity to the left visual cortex, to support the headplate with dental cement (Kemdent, UK) and Kwik-cast (WPI, UK). Over the same surgical window, viral injections were performed in different locations for Tsc1-floxed and C57BL/6J mice. In the Tsc1-floxed animals, burr-holes were drilled at the following coordinates relative to bregma: ML, +2.0 mm; AP, -2.6 mm; DV, 600 µm and 350 µm. To focally knockout floxed Tsc1 in astrocytes, we performed a high titer (1 x 1013 µg/ml) viral injection of AAV5- GFAP105-mCherry-Cre (AddGene, USA) at a rate of 75 nL/min (microinjection pump, WPI, USA). The volume at each respective depth was 400 nL and 200 nL, repeated at both sites. Delivery was achieved through a 5 µl syringe (Mo.5, Hamilton, Switzerland) in the somatosensory/visual cortex. In the CSD modulation animals, equipment remained constant, but injection sites, volumes and rates were varied. Two locations at the M2 area were selected relative to bregma: (1) ML, +0.8 mm; AP, +1.4 mm; DV, 600 µm and 350 µm, and (2) ML, +1.0 mm; AP, +2.3 mm; DV 600 µm and 350 µm. Injected volumes were 250 nL and 250 nL at each depth and site, at a rate of 50 nL/min. A high titer (1 x 1013 vg/ml) AAV9-CamK2a-ChR2-EYFP design was selected to express ChR2 locally. Animals were habituated in the head-fixed Neurotar system for periods of 15, 30, and 60 minutes, prior to experiments. To place the hybrid array on the brain, a craniotomy (2x2 mm) was performed on the same day of experiment over the somatosensory cortex. Additionally, to place the refences for the transistors and electrodes, we drilled two holes in the contralateral hemisphere (motor cortex for electrode reference – Pt; somatosensory for transistor reference – Ag/AgCl). Finally, in the case of CSD modulation experiments, we exposed the dura in the ipsilateral motor cortex to enable optogenetic stimulation through an optic cannula. Animals were left to recover 3-6 hours post-surgery before experiments began.

### In vivo recordings

Animals were head-fixed during the experiments in the previously habituated platform and the device was positioned over the somatosensory cortex through micromanipulators. The device was connected to the corresponding hardware through a custom printed circuit board. In the CSD modulation experiments, an LED cannula (200 µm diameter, Thor Labs, USA) was positioned over the ipsilateral motor cortex. The optic stimulus was a 10-second light-on (470 nm) protocol implemented in the CLD 1015 Thor Labs controller and externally modulated from Spike2 software (version 9.16; RRID:SCR_000903). The hardware and software of the setup were similar to in vitro experiments and consisted of a series of generic and custom-built devices and programs, respectively. To acquire signals through the gSGFETs, a custom-built 16-channel amplifier (sampling frequency: 9600 Hz; g.RAPHENE, g.tec GmbH, Austria) with large dynamic range was employed. Data was visualized in real time through a Simulink model (MATLAB 2016b RRID: SCR_001622, g.tec high speed library). To record through the glass micropipette, a Data Acquisition Unit (1401, Cambridge Electronic Design), and a 700B Multiclamp amplifier (sampling frequency: 10000 Hz) was used with Spike2 software. Electric stimulation was configured in MC_stimulus II (version 3.5.11) and delivered by the MCS stimulus generator STG4008 (8- channel) and STG4002 (2-channel; Multichannel systems, Harvard Bioscience Inc.). A subset of experiments was carried out with the RHS Stim/Recording Controller (Intan Technologies) and 16-channel RHS Intan head stages.

## Acknowledgements

This research was funded by the European Union’s Horizon 2020 research and innovation programme under grant agreement no. 881603 (Graphene Flagship Core 3) and the European Union’s Horizon Europe research and innovation programme under grant agreement 101070865 (MINIGRAPH) and 101130650 (META-BRAIN). The ICN2 has been supported by the Severo Ochoa Centres of Excellence programme [SEV-2017-0706]. ICN2 and IMB-CNM are currently supported by the Severo Ochoa Centres of Excellence and Maria de Maeztu Units of Excellence programme, grant CEX2021-001214-S and grant CEX2023- 001397-M, respectively, both funded by MCIN and MCIU/AEI/10.13039.501100011033. ICN2 is also supported by the CERCA Programme of Generalitat de Catalunya. This research is also funded by the Spanish Ministerio de Ciencia e Innovación (PID2020-113663RB-I00 and PID2021-126117NA-I00) founded by MCIU/AEI/10.13039/ 501100011033) and by “ERDF A way of making Europe”, PLEC2022-009232, PCI2021-122095-2A and CNS2023-144492, funded by MCIU/AEI /10.13039/501100011033 and the European Union Next-GenerationEU/PRTR. N. R. acknowledges grant PRE2020-093708 founded by MCIN/AEI /10.13039/501100011033 and by FSE. E. M. C. acknowledges grant FJC2021-046601-I funded by Agencia Estatal de Investigacion of Spain and the European Union Next-GenerationEU/PRTR. This work has made use of the SpanishICTS Network MICRONANOFABS, partially supported by MICINN and the ICTS NANBIOSIS, specifically by the Micro-NanoTechnology Unit U8 of the CIBER-BBN. This research was supported by CIBER - Consorcio Centro de Investigación Biomédica en Red- (CB06/01/0049), Instituto de Salud Carlos III, Ministerio de Ciencia e Innovación. The project that gave rise to these results received the support of a fellowship from the “la Caixa” Foundation (ID 100010434). M. P. acknowledges grant with fellowship code LCF/BQ/DI21/11860021.

## Contributions

1. M. P. worked on the design, fabrication, characterization and development of the devices; worked on the *in vivo* experimental design, data collection, and analysis of the *in vitro* data and *in vivo* data; worked on the preparation of figures and writing of the manuscript, participated in and contributed to the in vivo experiments.
2. M. E.-I. worked on the *in vivo* experimental design, data collection, and analysis; contributed to animal handling; carried out histological sectioning and imaging; preparation figures and writing of the manuscript E. M.-C contributed to the characterization of the devices, data analysis, and manuscript preparation.
3. X. I. and N. R. contributed to fabrication.
4. N. C. and D. R. carried out animal surgeries, habituations, and in *vivo* experiments.
5. K. K., E. C. and R. G.-C contributed to the manuscript preparation.
6. R. W., A. G.-B. and J. A. G. designed and supervised the project and contributed to the manuscript preparation.

